# SHARE-Topic: Bayesian Interpretable Modelling of Single-Cell Multi-Omic Data

**DOI:** 10.1101/2023.02.02.526696

**Authors:** Nour El Kazwini, Guido Sanguinetti

**Affiliations:** Theoretical and Scientific Data Science, Scuola Internazionale Superiore di Studi Avanzati, Trieste, Italy

**Keywords:** Gene regulation, Single-cell multi-omics, Bayesian modelling, Interpretability, Gene regulator in cancer, Lymphoma

## Abstract

Single-cell sequencing technologies are providing unprecedented insights into the molecular biology of individual cells. More recently, multi-omic technologies have emerged which can simultaneously measure gene expression and the epigenomic state of the same cell, holding the promise to unlock our understanding of the epigenetic mechanisms of gene regulation. However, the sparsity and noisy nature of the data pose fundamental statistical challenges which hinder our ability to extract biological knowledge from these complex data sets. Here we propose SHARE-Topic, a Bayesian generative model of multi-omic single cell data which addresses these challenges from the point of view of topic models. SHARE-Topic identifies common patterns of co-variation between different ‘omic layers, providing interpretable explanations for the complexity of the data. Tested on joint ATAC and expression data, SHARE-Topic was able to provide low dimensional representations that recapitulate known biology, and to define in a principled way associations between genes and distal regulators in individual cells. We illustrate SHARE-Topic in a case study of B-cell lymphoma, studying the usage of alternative promoters in the regulation of the FOXP1 transcription factors.

## Introduction

Biological complexity arises from interactions of many molecular factors at varying spatial and temporal scales. Understanding the nature and dynamics of these interactions is a major open problem in fundamental biology, with potentially important translational implications. Over the last two decades, the emergence of next generation sequencing technologies, and more recently of single-cell sequencing technologies, has been a major accelerator towards tackling these questions, with large international consortia such as ENCODE and the Human Cell Atlas (1, 2) providing the community with invaluable data sets measuring a variety of molecular features potentially influencing gene expression.

Recent breakthroughs in single-cell technology have opened the possibility of measuring, in a high-throughput fashion, multiple molecular layers within the same cell, providing new opportunities to enhance our understanding of the interactions between biological factors. These technologies, collectively referred to as single-cell (sc) multi-omics, are generally designed to measure simultaneously the cell’s transcriptome together with one or more other molecular features, typically epigenetic factors such as DNA methylation or chromatin accessibility, DNA sequence or presence of protein markers (3–6). These sc multi-omics technologies offer in principle a number of enticing possibilities, chief among them the opportunity to measure gene regulatory mechanisms in individual cells, and its variability across cells.

In practice, analysing and interpreting sc multi-omic data present considerable challenges, due to the high level of sparsity and noisiness of the data (7). To date, most efforts have focussed on deploying different deep learning architectures within the variational autoencoder (VAE) paradigm (8–12). VAEs use non-linear maps (parametrised by deep neural networks) to explain the variability in the data in terms of a latent space, where each cell is represented by a low dimensional vector (typically of around ten dimensions, instead of tens of thousands of molecular features). Analysis of the resulting latent spaces have shown to outperform basic dimensional reduction methods in terms of clustering cells and for survival analysis (8–10). Despite these successes in extracting patterns at the cell level, obtaining insights at the gene level from VAEs (for example in terms of specific regulatory interactions) is extremely difficult, due to the effective impossibility of reliably interpret the contribution of individual genes in complex nonlinear models. Indeed, even simple linear analyses, such as measuring correlations between region accessibility and gene expression, are very challenging in the single-cell realm due to the high levels of noise, as demonstrated recently in (13).

Here, we introduce SHARE-Topic, a Bayesian statistical model of joint chromatin accessibility and transcriptomic data, perhaps the most widely available type of sc multi-omic data. SHARE-Topic extends the cisTopic model of singlecell chromatin accessibility (14) by coupling the epigenomic state with gene expression through latent variables (topics) which are associated to regions and genes within an individual cell. In this way, SHARE-Topic is able to extract a latent space representation of each cell informed by both the epigenome and the transciptome, but crucially also to model the joint variability of individual genes regions, providing an interpretable analysis tool which can help in generating novel hypotheses from the data. We test SHARE-topic on two sc multi-omics data produced from SHARE-seq protocol, a small data set (~ 3000 cells) coming from mouse brain and a relatively bigger data set (~30000 cells) of mouse skin. The model demonstrates good scalability of the algorithm as well as its ability to extract novel biological information from these complex data sets. As a more biologically compelling example, we then analyse a recent lymphoma data set generated using the commercial 10X multiome technology. We investigate the use of alternative promoters in the regulation of the master regulator FOXP1 in tumour cells, a putative tumour suppressor whose complex transcriptional architecture was already associated with B-cell lymphoma (15, 16). We show a clear preference for the usage of one specific internal promoter, providing a case study of how non-trivial biological information can be derived from analysing these rich data sets.

## Results

### The SHARE-Topic model

Topic models are unsupervised learning algorithms, originally designed to analyze and annotate large archives of text documents with thematic information (17–21). The premise of topic modelling is that each document can be represented as a point in a much lower dimensional space (topic space), corresponding to the relative importance of different topics to the document. The probability of a word appearing in a document depends strongly on the topic, hence each topic is associated with a distinct distribution over word frequencies, which can also be used to associate topics with semantically meaningful annotations.

Bravo González-Blas et al. (14) have recently proposed cis-Topic, a topic model designed to efficiently analyse single-cell ATAC-seq data. cis-Topic provides an effective tool to obtain lower dimensional representations of the very high-dimensional scATAC-seq data, however interpretation of its latent space is complicated by the varying quality of the annotation of open chromatin regions. In this paper, we present SHARE-Topic, a topic model adapted to multi-omics data which allows both a stronger interpretability and genelevel predictions. A high-level view of the model structure is given in Figure 1a: single-cell multi-omics, encoded as two high-dimensional sparse matrices, is the input to SHARE-topic. The model than utilises a Gibbs sampler to obtain posterior estimates of the various parameters, which can be used both to obtain a low dimensional representation of the cells, and to associate topics to cells and regions to genes. The structure of the model is given in Figure 1b. A table illustrating the correspondence of concepts in classical (text based) topic modelling and their multi-omics analogue in SHARE-topic is given in Table 1.

**Fig. 1.**
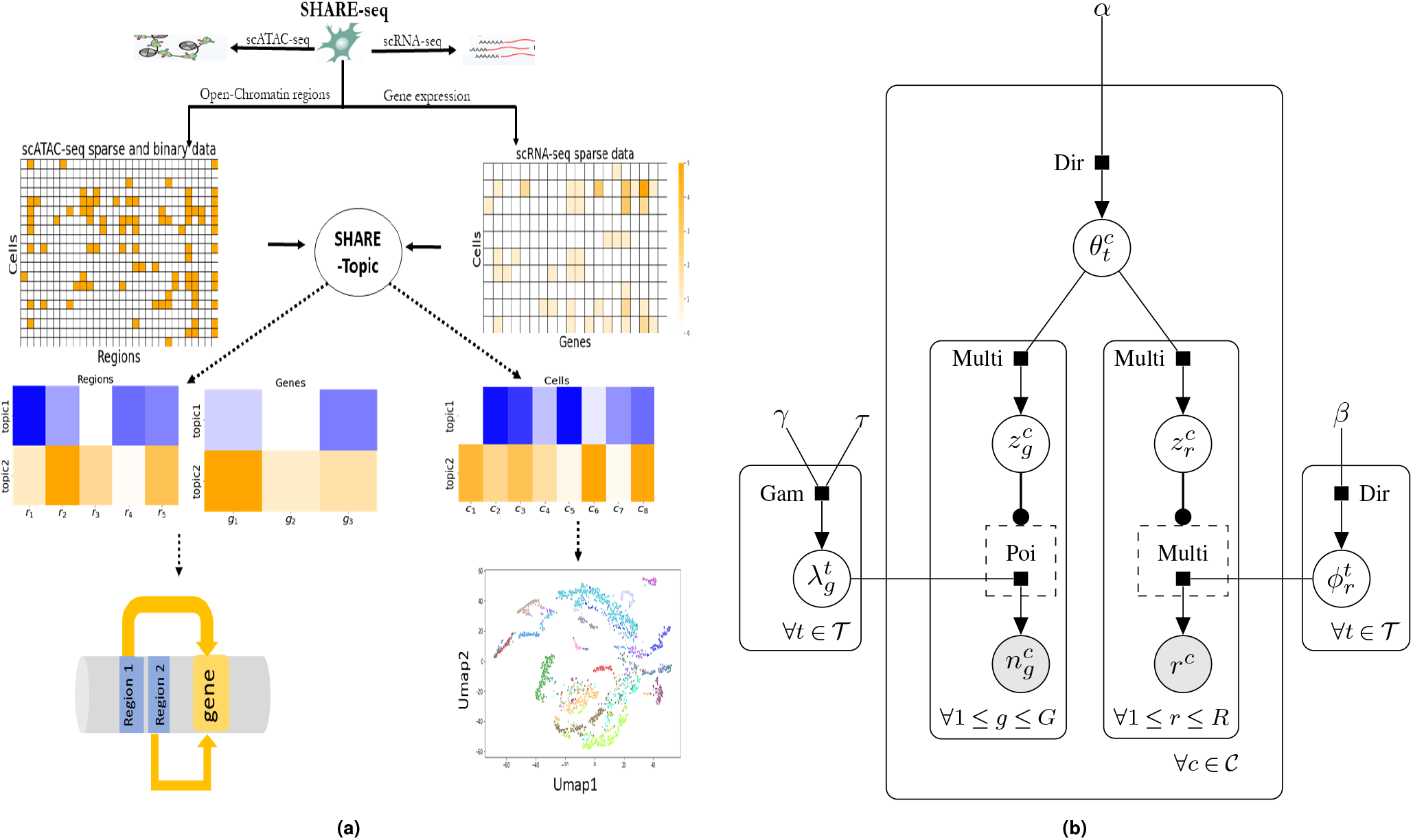
(a)The general workflow for SHARE-Topic. Share-Topic is an extended LDA(Gibbs sampler) model to accommodate both scRNA-seq and scATAC-seq data (multiomics) provided by SHARE-seq. Share-Topic cluster cells and infer associate regions with genes.(b)The graphical model of SHARE-Topic.

**Table 1.**
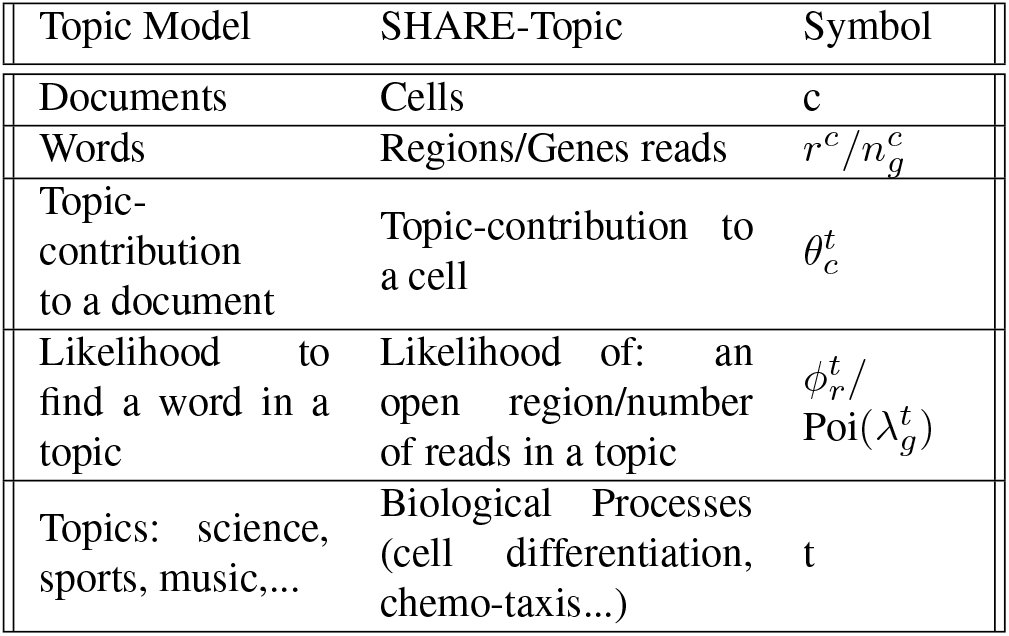
Interpolation of topic model to biological framework.

### Data sets used

We use two data sets generated with the SHARE-seq technology (6), as well as one data set generated with the commercial 10X multiome platform. The SHARE-seq data sets profile approximately 3 thousand mouse brain cells and approximately 30 thousand mouse skin cells; the number of genes/ regions retained for each data set after pre-processing is of approximately six thousand genes expressed, and 2 × 10^6^ regions for the brain data set, and approximately 3100 genes, and 9 × 10^5^ regions for the skin data set. The multiome data set profiles approximately 14 thousand lymphoma cancer cells, retaining approximately 8 thousand genes and 1.2×10^4^ regions. Details of the filtering procedure can be found in the Methods section.

### SHARE-Topic recapitulates cell identities

As with any other dimensionality reduction tool, from PCA to variational auto-encoders, a primary output of SHARE-topic is the assignment of a latent vector to every cell. In our case, this vector is a probability distribution over the topics indicating to which topic each cell partakes. The choice of the number of topics (dimensionality of the latent space) is a nontrivial hyperparameter tuning issue; we resort to using the Watanabe Information Criterion (WAIC) (22) (more details on the criteria for the choice of topics number are given in the method section). The latent space can then be visualised using tools such as Uniform Manifold Approximation and Projection (UMAP) (23), and the consistency of the visualisation with existing annotations can be assessed using quantitative criteria.

Figure 2 shows the results of this exercise on the three data sets we consider. The panels show a UMAP reduction to two dimensions of the (posterior mean) topic vectors assigned to each cell, with each dot coloured according to the corresponding cell-type annotation. Visually, all three plots highlight a good separation between cell types and a biologically plausible organisation of the latent space.

**Fig. 2.**
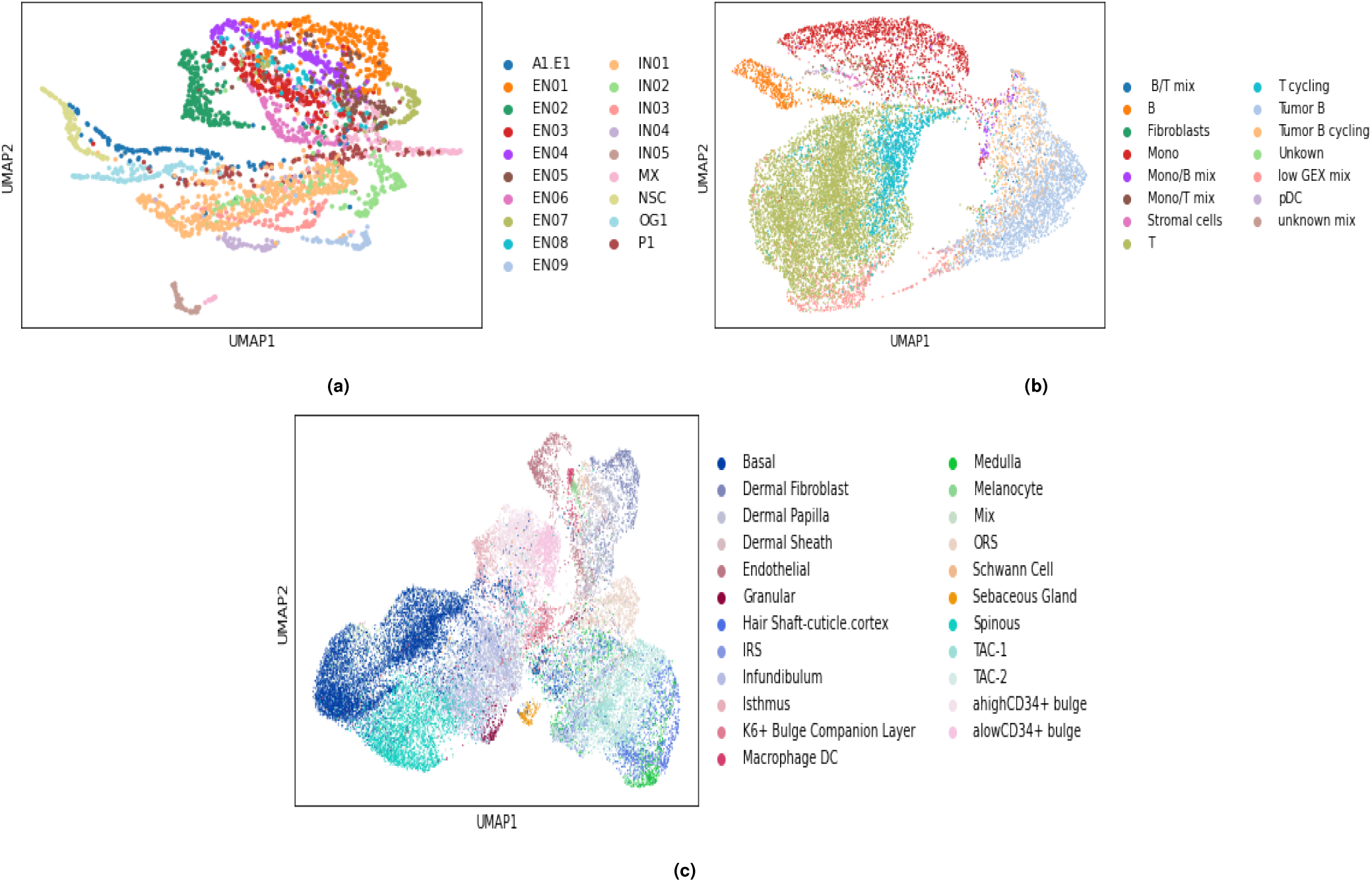
UMAP embedding of SHARE-Topic based on cell-topic distribution(*θ^c^*). (a) brain data set embedding of 2781 cells from topic space of dimension 30. (b) B-cells lymphoma data set embedding of 14566 cells from topic space of dimension 45. (c) skin data set embedding of 27782 cells from topic space of dimension 60.

Naturally, SHARE-topic is not the only method capable of obtaining a latent representation from multi-omic data. To compare SHARE-Topic with the state-of-the-art, we performed dimensionality reduction using Multi-Omic Factor Analysis (MOFA+, (24)) a recently proposed linear dimensionality reduction method specifically taylored for singlecell multi-omic data. While MOFA+ is potentially less flexible than nonlinear alternatives such as VAEs, its linear nature makes is strongly interpretable, thus making it a fair compar ison with SHARE-Topic. We used the implementation within the muon platform (25); due to a technical problem, related to memory usage, we couldn’t run MOFA+ on the whole mouse skin and lymphoma datasets and therefore subsampled that data set retaining approximately only 25k chromatin regions for the ATAC component.

To benchmark the quality of the data integration performed by SHARE-Topic, we also compared the performance of SHARE-Topic with the reduced models obtained by considering only the transcriptome or the chromatin accessibility data (right and left arms of the graphical model in Figure 1b). The corresponding visualisations are given in Supplementary Figures S4 and S5.

To assess quantitatively the validity of the latent representation discovered by SHARE-topic, we use a *k*-Nearest Neighbour (*k*-NN, using *k* = 10) classifier trained on 70% of the cell type annotations, and evaluate its accuracy in predicting the annotations of the remaining 30% of cells. The results of this assessment are shown in Table 2, and in Figure S3 in terms of confusion matrices. For both SHARE-seq data sets, all methods perform significantly better than random (which would have an accuracy of around 5% since there are around 20 cell types in all data sets), indicating that the single-cell multi-omic data captures salient features of cell identity. In both data sets, SHARE-Topic outperforms MOFA+ by a very large margin, suggesting that the topic representation is indeed better reflecting cell identity when compared with the generalised linear structure of MOFA+. SHARE-topic also significantly outperforms its RNA-only or accessibility-only version, indicating that the model is able to effectively integrate both channels of information. This is particularly marked in the brain data set, where we also notice a significant difference in the accuracy of the RNA-based and ATAC-based classifiers. For the skin data set, both sources independently achieve a good level of accuracy, however the joint model still outperforms both. Thus, we conclude that both sources carry valuable and, to some extent, complementary information towards classifying the cells.

**Table 2.**
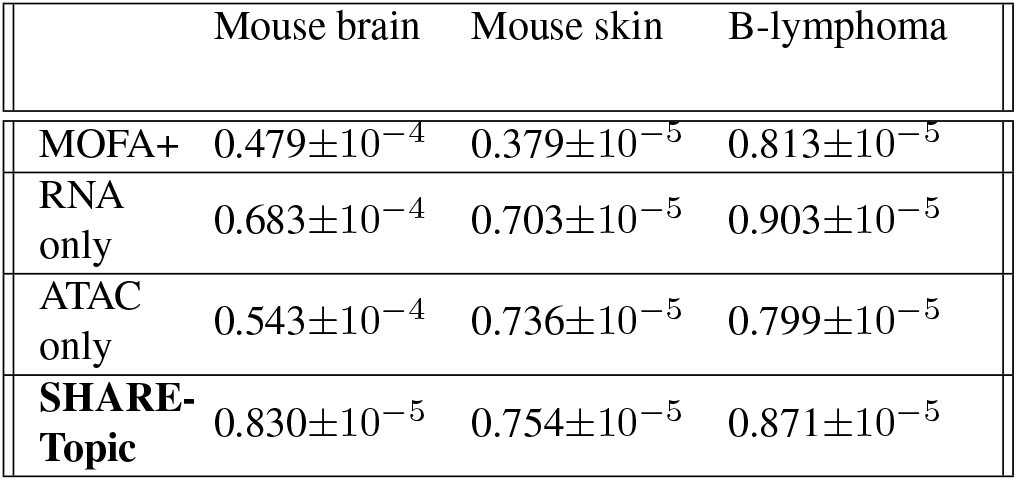
Table showing accuracy of *K*-NN classifiers trained on the latent representation of different methods to predict cell types in the three datasets.

The story is slightly different for the lymphoma multiome data set. Here we have a much stronger performance by MOFA+, although still considerably below the level of SHARE-Topic. What is different is however that inclusion of the ATAC-seq modality seems to slightly degrade the performance of the topic model, which is in fact best run on RNA-seq only. We do not know whether this issue is related to the technology used, the biology of the lymphoma cancer cells, or to the way the cell type annotations were derived.

### Associating SHARE-Topic results to underlying biology

SHARE-Topic’s performance at identifying effective low dimensional representation supports our hypothesis that the degrees of freedom of the system are far fewer than the dimensions of the very high dimensional spaces of genes and regulatory regions. This hypothesis is shared by all dimensionality reduction approaches developed for single-cell multi-omics. SHARE-Topic, due to its transparent probabilistic formulation, offers a natural way to interpret its results, making it suitable as a hypothesis-generating tool.

One simple approach to interpret SHARE-Topic results is to consider topic assignments at the cell level. For example, one may select all cells with the same dominant topic (largest element of the cell-topic assignment vector *θ_c_*) and then check for enrichment of specific cell types among the selected set.

Alternatively, one may leverage the gene by topic matrix, whose entries 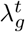 provide expected expression levels of a gene in a certain topic, to obtain a molecular interpretation of the SHARE-Topic latent space in terms of biological processes associated with each topic. To do so, we first associate genes to a topic by computing an entropic measure of the distribution of gene expression across topics (Figure 3a, see Methods). Intuitively, we seek genes which are highly expressed in one (or few) topics, and nearly silent in the others; such genes will have a very low entropy, indicating a distribution across topics which is far from uniform (Figure 3b). By pre-filtering genes based on low entropy levels, we can then associate each gene to the top topic in terms of corresponding expression. This procedure enables us to associate a set of genes to each topic, which can then be queried for enrichment using tools such as clusterProfiler(26, 27).

**Fig. 3.**
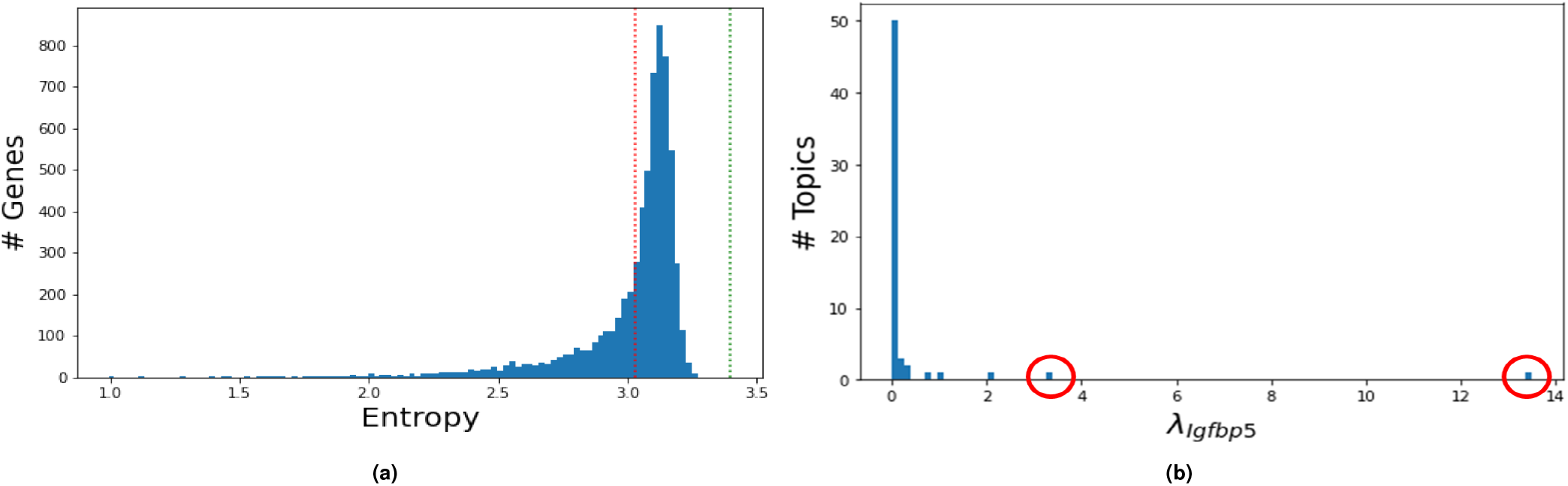
(a)Histogram of the entropy distribution of expression rates across the 30 topics in the brain data set (6421 genes). The green line corresponds to the entropy of a uniform distribution (Bernoulli(1/30)). A gene with strong membership to a certain topic will have *λ*s with low entropy across topics, indicating a highly non-uniform distribution over the topics. The red line represents the 30 percentile of genes that we consider to have a clear membership to a topic and we include them in the downstream analysis. (b) Histogram for distribution of the average expression of example gene Igfbp5 (*λ_Igfbp_*_5_) across topics. The red circles show that there are topics with high average expression compared to other topics that have average expression around zero. This suggest that this gene has strong membership to these topics and thus contributes significantly to the biological process associated with it.

An example of these analyses is provided in Figure 4 for the brain data set. Here a particular topic (topic 27 in our run of the algorithm) was strongly concentrated within a particular cell type, oligodendrocytes (as shown in the UMAP visualisation in Fig 4a). Considering genes significantly associated with the topic, a number of enriched gene functions appear, primarily but not solely connected with neural function and development. A table summarizing the principal functions associated with the main topics in each data set is provided in the Supplementary Table S1.

**Fig. 4.**
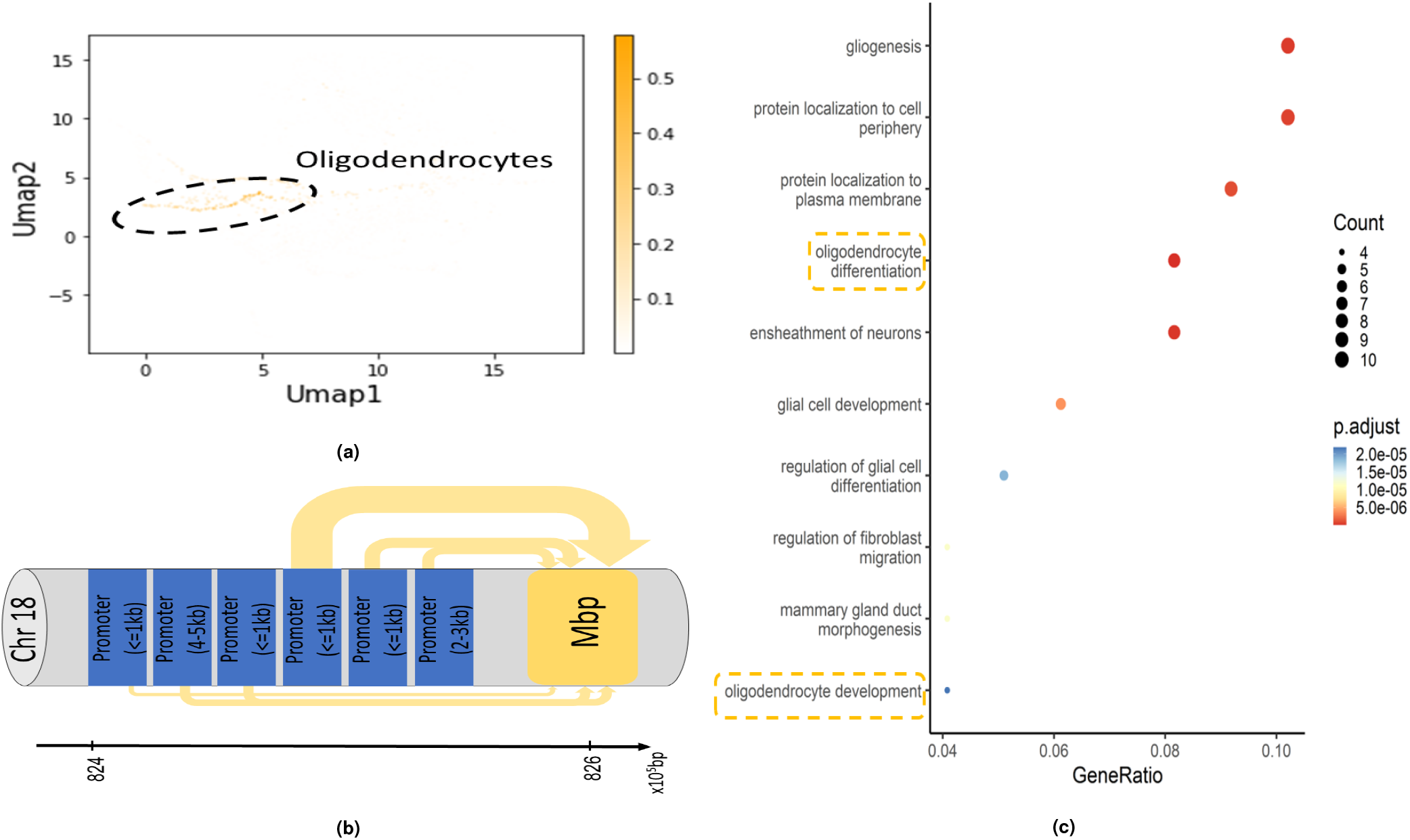
Analysis of topic 27 in the brain data set. There are 9192 regions and 107 genes associated with topic 27. (a) Umap embedding of brain data showing the topic 27 contribution to cells. This topic is enriched in a specific type of cell (Oligodendrocytes). (b) The interaction probability between annotated promoters and the Mbp gene(myelin basic protein) conditioned on membership of topic 27.(c) The biological processes enriched in topic 27. Two biological processes are oligodendrocytes specific.

### SHARE-Topic uncovers regulatory events

One of the most attractive features of sc multi-omic data is the opportunity to identify cell-specific gene-region associations. To do this, we identify pairs of genes and chromatin regions (within a certain neighbourhood) which have a strong joint probability of expression for the gene and of opening for the region, conditioned on the topic. In practice, we take a gene-centric view: we start with genes strongly associated to a topic. We then select annotated regions in a pre-defined neighborhood of 105bp using the CHIPseeker package (28, 29), and rank regions according to their probability of being open in the gene-associated topic. The choice of a very large window of 10^5^bp is designed to capture both distal and proximal regulatory relationships.

An example of the outcomes of this type of analysis is shown in Figure 4). The figure focuses on the MBP (myelin basic protein) gene, which codes for a protein which is essential in the development of the myelin sheath encasing oligodendrocytes. Its expression is strongly associated with the oligodendrocyte topic, consistently with its annotation, and can be driven by as many as six distinct annotated promoters, three of which in the proximal region (less than 1kb away). Figure 4b shows the open probability of each of these six regions (in the oligodendrocyte topic) as inferred by SHARE-Topic in the brain data set, depicted as the thickness of the arrows connecting the promoter region with its target genes. As one of the arrows dominates the others, this analysis suggests that transcription is primarily driven by one of these six promoters (schematically represented as the third from the right). Similar gene-card type diagrams are shown for other genes in the two SHARE-seq data sets are provided in Supplementary Figure S6.

### SHARE-Topic elucidates the regulation of FOXP1 in B– cell lymphoma

As a more biologically compelling example of the use of SHARE-Topic, we turn to the lymphoma multiome data set and perform a similar analysis. We focus on topic 13; this topic is primarily associated with B-cell tumour cells, and presents a strong enrichment of the cytokine production pathway, indicating a probable involvement in the inflammatory response.

Among the prominent genes associated with this topic, we focus on the master regulator FOXP1, an essential gene in development which has been associated with several cancers, including lymphoma (16). FOXP1 is a long gene (approximately 600Kb) with a complex transcriptional architecture, expressing several isoform; excess abundance of a short isoform has been reported to be a marker for lymphoma (15). Brown et al. (15) also observed the presence of two predicted internal promoters just before the start of the short isoform. In our data set, we find that open chromatin regions are measured for three possible promoters; Figure 5b shows the relative importance of the three promoters conditioned on the cells being in topic 13. Once again we notice a clear dominance of one promoter over the others; while this is a purely correlative observation at this level, it represents another example of the type of non-trivial testable predictions that can be produced by SHARE-Topic.

**Fig. 5.**
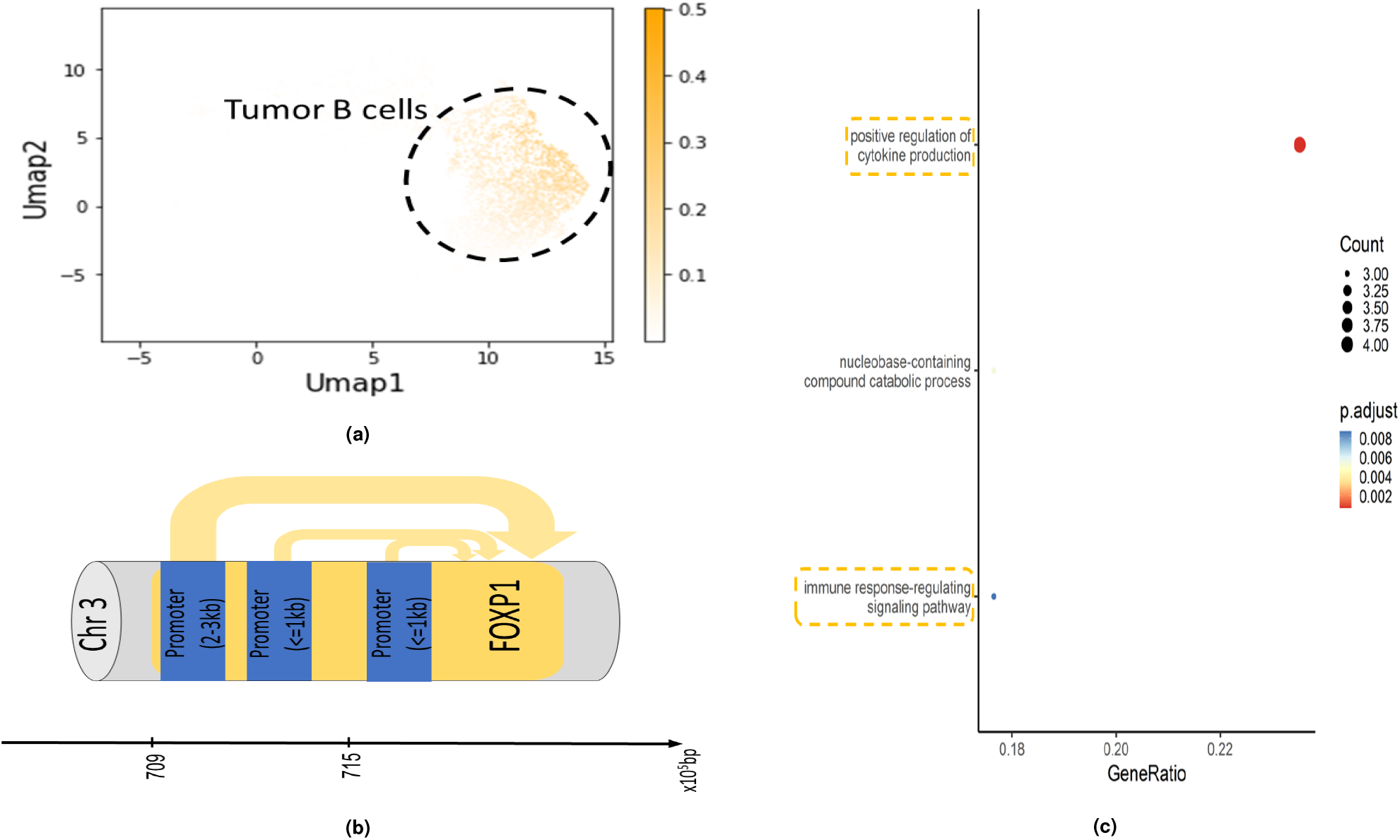
Analysis of topic 13 in the lymphoma data set. There are 2031 regions and 24 genes associated with topic 13.(a) Umap embedding of lymphoma data showing the topic 13 contribution to cells. This topic is enriched in tumor B cells. (b) The interaction probability between promoters and FOXP1 gene associated with the same topic. (c)The biological processes enriched in topic 13.

### Conclusions

Single-cell multi-omic technologies open up unprecedented opportunities to explore the molecular landscape of living cells. Most existing deep-learning based methods have focussed on representing the variation in these data sets at the cell level, developing tools which focus on highlighting the diversity across cell populations, but often at the cost of hiding in algorithmic complexity the molecular mechanisms which give rise to this diversity.

With SHARE-Topic, we propose a Bayesian hierarchical model with transparent probabilistic semantics for the analysis of joint expression and chromatin accessibility data. SHARE-Topic provides a low-dimensional representation of multi-omic data by embedding cells in a topic space. We show on two data sets that SHARE-Topic embeddings are highly accurate at recapitulating cell diversity and effectively integrate both channels of information. Moreover, the simplicity of the model enables a straightforward interpretation of the obtained embeddings in terms of biological processes and permits non-trivial gene-level insights on the interactions of chromatin accessibility and gene expression in single cells.

SHARE-Topic can also be extended to build more complex data models which will inevitably be needed as singlecell multi-omics technologies become more widely used in biomedicine. In particular, the availability of commercial kits, such as the 10X multiome for simultaneous single-cell expression and chromatin profiling, will require the development of models which can dissect variability arising from multiple donors and multiple (possibly related) conditions, creating new challenges for method development.

## Methods

### Data filtering

In order to remove noise from the data we looked first at the transcriptome profile of cells and removed outlier cells. In each cell type, cells with a total number of genes read lower than 5-10% or higher than 90% percentile are considered an outlier and discarded in our study. In the lymphoma data set, all cells were retained due to the high heterogeneity of tumour cells. For the transcriptomic data, genes that are expressed in over 5% of the cells are considered.

Since the chromatin accessibility data is binary and highly sparse, filtering the regions is decided according to the average number of region reads across cells. For the skin and lymphoma data set, we retain regions present in at least ~ 1% of all cells. For the brain data set, which has lower coverage in ATAC-seq, the threshold is 10 fold less (~ 0.1%).

### SHARE-Topic implementation

SHARE-Topic is designed to derive from the multiome dataset (transcriptome and chromatin accessibility) a regulatory topic space of dimension T (number of topics). The implementation is based on LDA and extends to include multiple inputs to infer interaction between inputs in the reduced dimension topic space. SHARE-Topic infers:

1. - 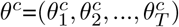: probability distribution of topics in a cell c. 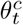 represents the contribution (importance) of a topic t to a cell c.
2. - 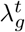: Poisson rate for gene reads in a topic. i.e the average number of expected reads when the gene is contributing to a topic t. The lambdas are considered independent in our model across topics and genes.
3. - 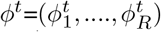: probability distribution of regions in a topic. 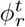 is the likelihood to observe a region r in a topic t.

SHARE-Topic is implemented using a Gibbs sampler and the update equations are derived based on the SHARE-Topic graphical model shown in 1b. The latent variables are initialized from predefined priors:

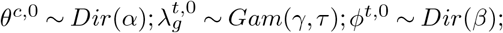

where:

*α, β*: pseudo-count for Dirichlet distribution
*γ, τ* : shape and scale parameters respectively of the gamma distribution

Using the conjugacy property between the priors and likelihood, the Gibbs update equations of the model at the *k*-the step are written as follows:

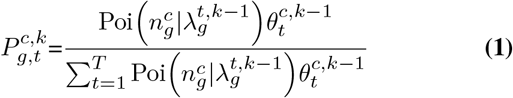

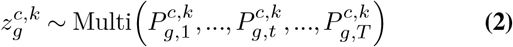

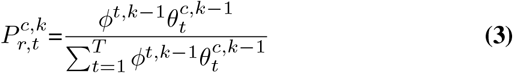

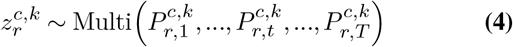

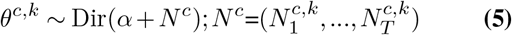

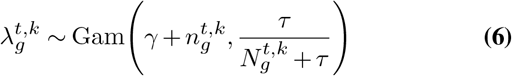

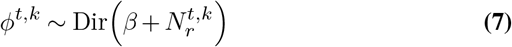

Such that:

- 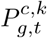: Probability of a gene g read in a cell c to have to have membership in a topic t,
- 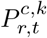: Probability of an observed region r in a cell c to have membership in a topic t,
- 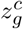:Topic membership of a gene g in cell c,
- 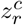:Topic membership of a region r in cell c,
- 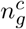:Count of a gene g in cell c,
- *r^c^*: Observed region r in cell c,
- 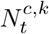: Total number of regions and genes that have t-th membership in cell c in the *k*-th step.
- 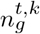:Total number of reads for gene g of t-th membership in the k-th step,
- 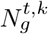:Total number of gene g across cells of t-th membership in the k-th step.
- 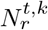:Total number of region r across cells of t-th membership in the k-th step,

The hyperparameters of the model are fixed such that *α* = 50/*T, β* = 0.1, *γ* = 1, and *τ* = 0.5. We run the MCMC to obtain 3000 samples where 500 samples are used as burn-in. To mild the effect of correlations, we considered a single sample every 10 samples. The convergence of the MCMC chain is assisted by monitoring the evolution of the likelihood (Figure S1).

The outputs of the Gibbs sampler are three matrices: 1) *θ* of dimension K× C× T, 2) *λ* of dimension K× T× G, 3) *ϕ* of dimension K× T× R. Such that K, T, C, and G are the number of samples, topics, cells, genes, and regions respectively. The latent parameters are estimated using the mean of the samples.

### Choosing number of topics

The number of topics is chosen according to the widely applicable information criterion(WAIC). Using the samples, WAIC of the model is obtained by computing the log-pointwisepredictive-density (lppd) and the variance in log probabilities for each observation (penalty term)(30):

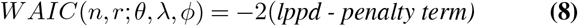

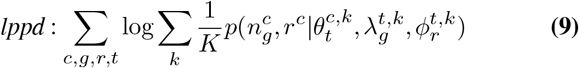

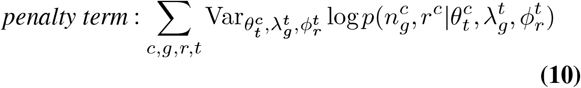

The WAIC is computed for both datasets for different numbers of topics (Figure S2). Based on that 30, 45, and 60 topics are chosen for brain, B-cell lymphoma, and skin datasets respectively.

### Associating genes with topics and topics annotation

Given the possibility that topics represent different biological processes, we expect genes to be strongly presented in certain topics and thus highly transcribed in a biological process while relatively less or absent in other topics. Capturing the genes with strong topic-membership means filtering out genes that have a non-uniform *λ*s distribution across topics (example: Figure 3b).

The relative contribution of a topic to a certain gene expression is calculated by dividing 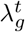 by the sum of the lambdas across topics and then used to compute the entropy of lambda for each gene. The distribution of the entropy for all genes is shown in Figure 3a and compared to the entropy value of an “event” that is uniform across topics. The genes with low entropy have lambdas that are relatively high in a few topics and almost vanish in the rest. The top 30% genes with the lowest entropy are considered to have a clear membership. For each of the selected genes, the top two topics with the highest lambdas are considered to be the topics associated with the gene.

After obtaining a group of genes that are associated with topics we carried gene ontology to determine the biological processes enriched in each of these groups and annotate the topics using a Bioconductor package, clusterProfiler (26, 27) (Table S1).

### Associating regions with topics

A similar procedure to topic-gene association is carried out on regions. The entropy of a specific region is computed across topics. Then top 30% of the regions with low entropy are considered to have a membership in specific topics. The selected regions are considered to be associated with two topics that have the maximum 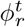 among other topics.

### Inferring regions-genes interactions

A Gene-region pair associated with the same topic simultaneously suggests a strong interaction within the pair. This interaction depends also on the genomic coordinates of the region and gene. Using the ChIPSeeker package in R the regions in the datasets are annotated (promoters(<=1kb), promoters(3-4 kb), intergenic distance,…), and the nearest gene to each region is returned also.

We looked at close region-gene pairs along the chromosome that are associated with the same topic, focusing on the regions that are annotated as promoters. Promoters that have higher 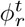 (more probable to be open) are considered to contribute more to the gene close to them.

## Data availability

The SHARE-seq data sets are publicly available on the GE0 website GSE140203, and B cell lymphoma data set is available on the 10xgenomics website here.

