## Supplemental materials for "SHARE-Topic: Bayesian Interpretable Modelling of Single-Cell Multi-Omic Data"

### 1 Appendix

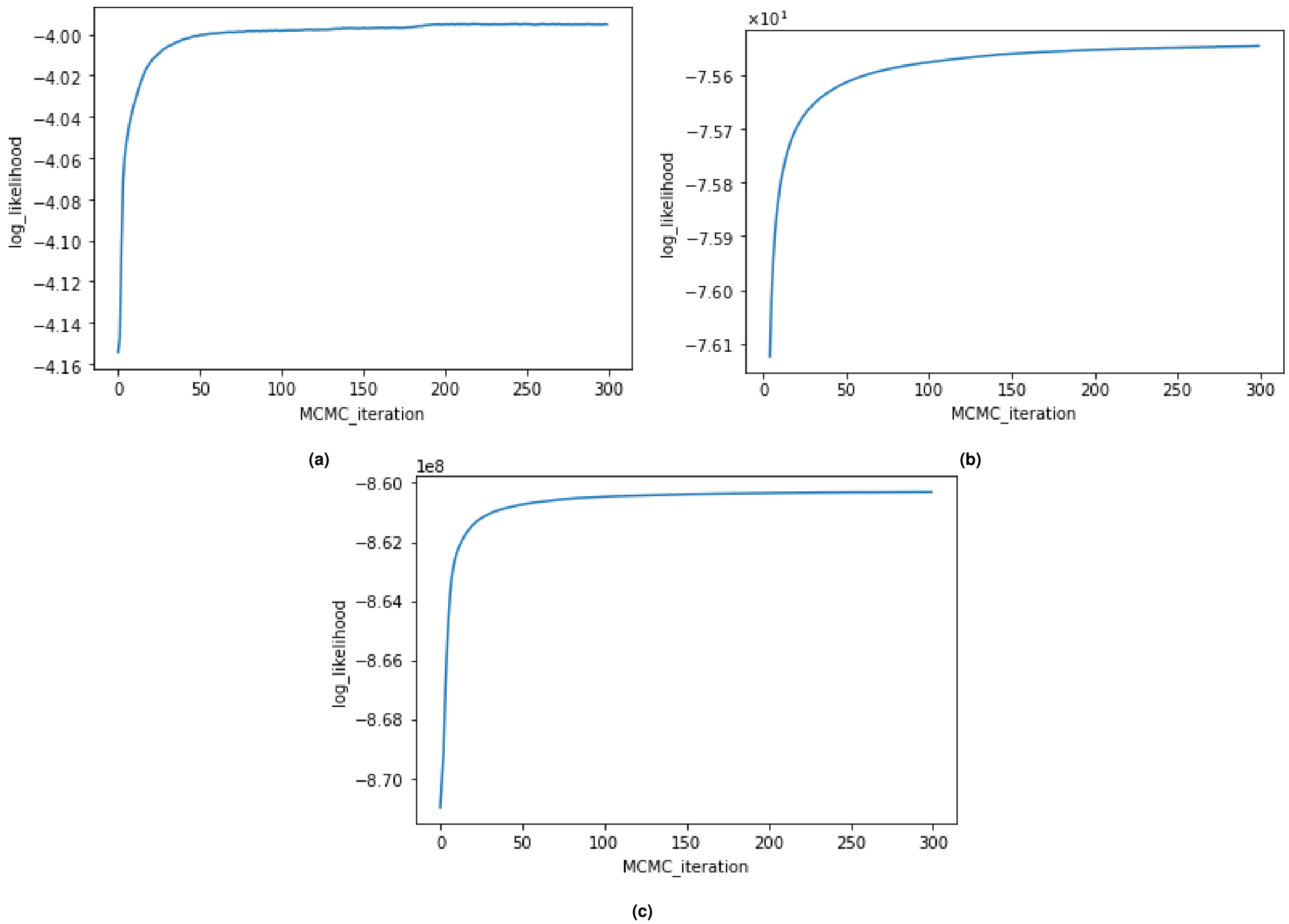


**Fig. S1.** Assessing convergence of the MCMC chains using the log-likelihood for: (a)mouse brain, (b) B-cell lymphoma, and (c) mouse skin datasets. The log-likelihood stabilizes after 50 samples(500 samples without the burnin).

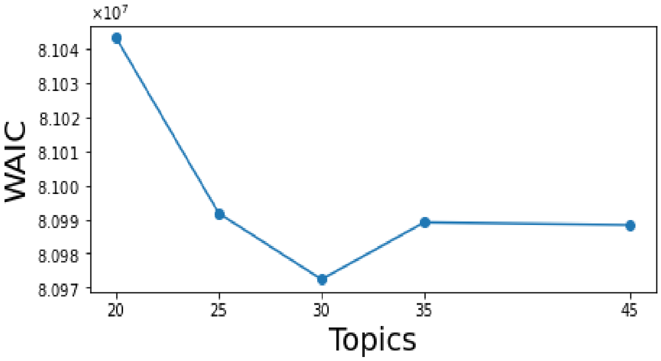


**(a)**

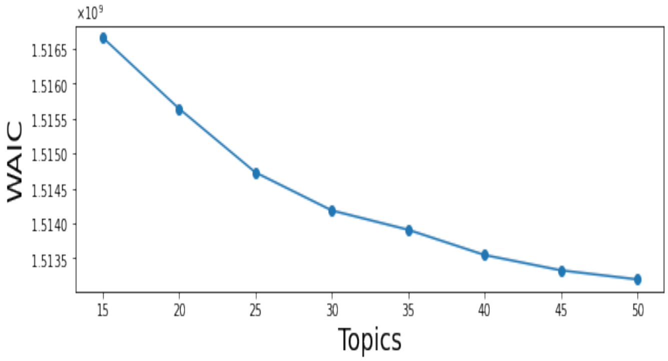


**(b)**

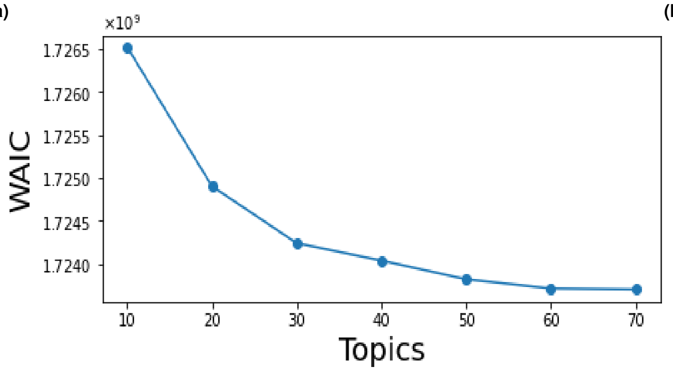


**(c)**

**Fig. S2.** Evolving of WAIC with number of topics. (a) For the brain datastet a minimum is reached at 30 topics. (b) for the B lymphoma dataset it is 45. (c) for skin dataset we choose 60 topics. For figures b and c we choose the number of topics such curve become relatively not sensitive when increasing topics.

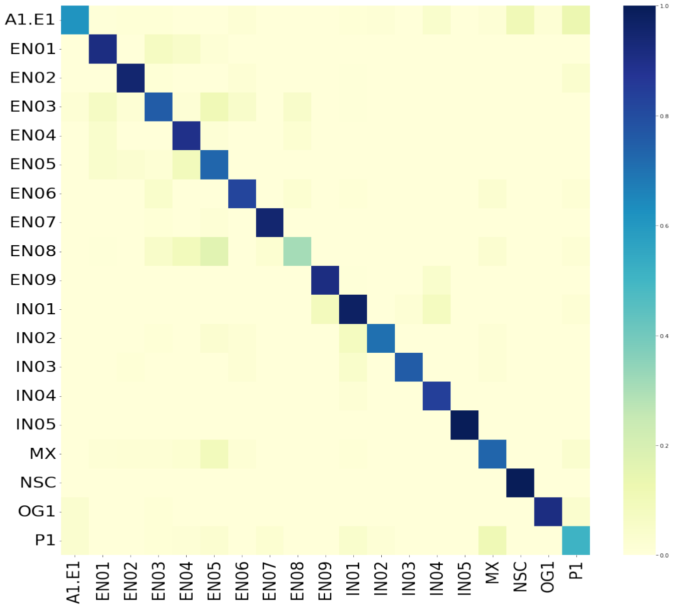


(a)

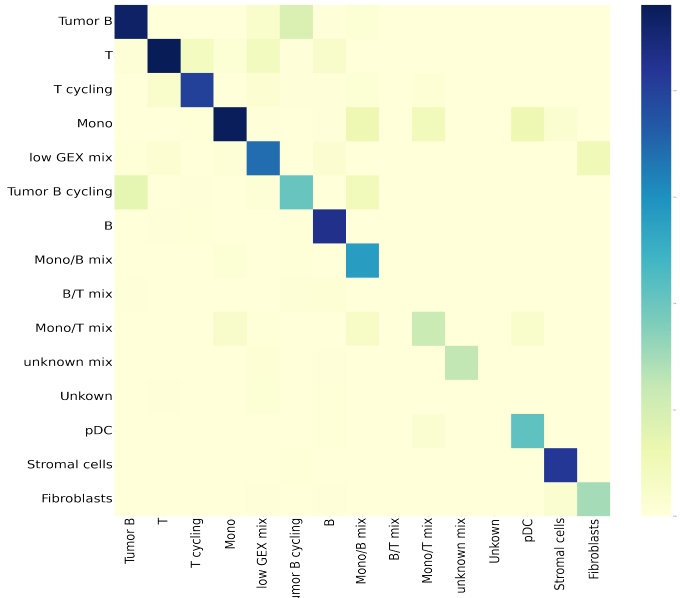


(b)

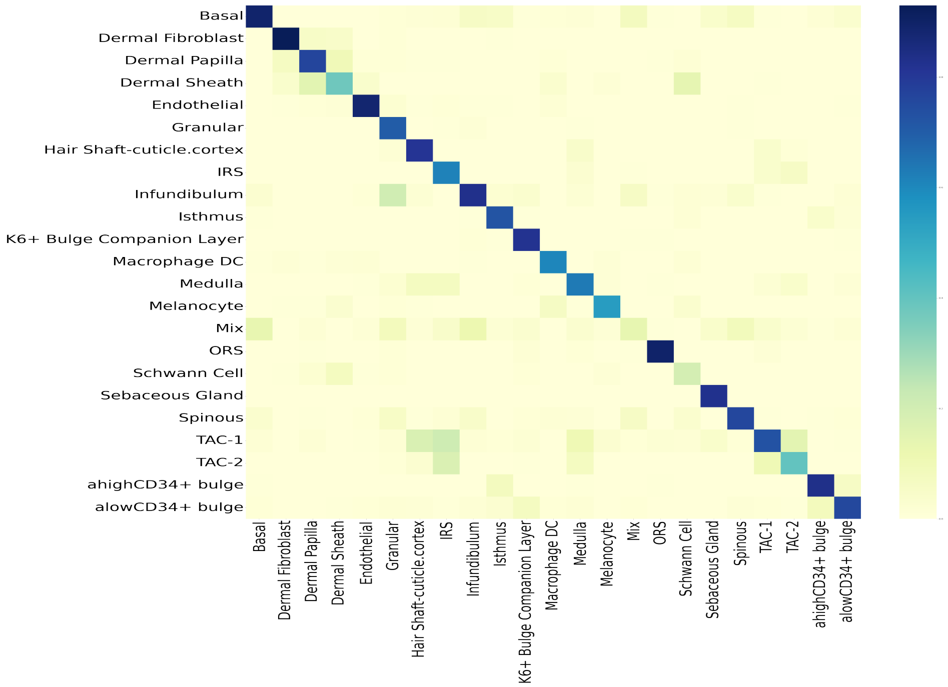


(c)

**Fig. S3.** Heatmaps of the confusion matrices showing the accuracy of KNN classifier in identifying different cell types. (a) In the brain dataset the classifier have good accuracy in identifying all types except for the case of EN08 it reaches the lowest accuracy 20%. (b) For the B lymphoma dataset the KNN do well in identifying strongly presented cell types and fails with the low presented cell population (Mono/B mix).(c) In the skin dataset the accuracy is relatively low for two types (ORS, Melanocyte) aslso 20%.

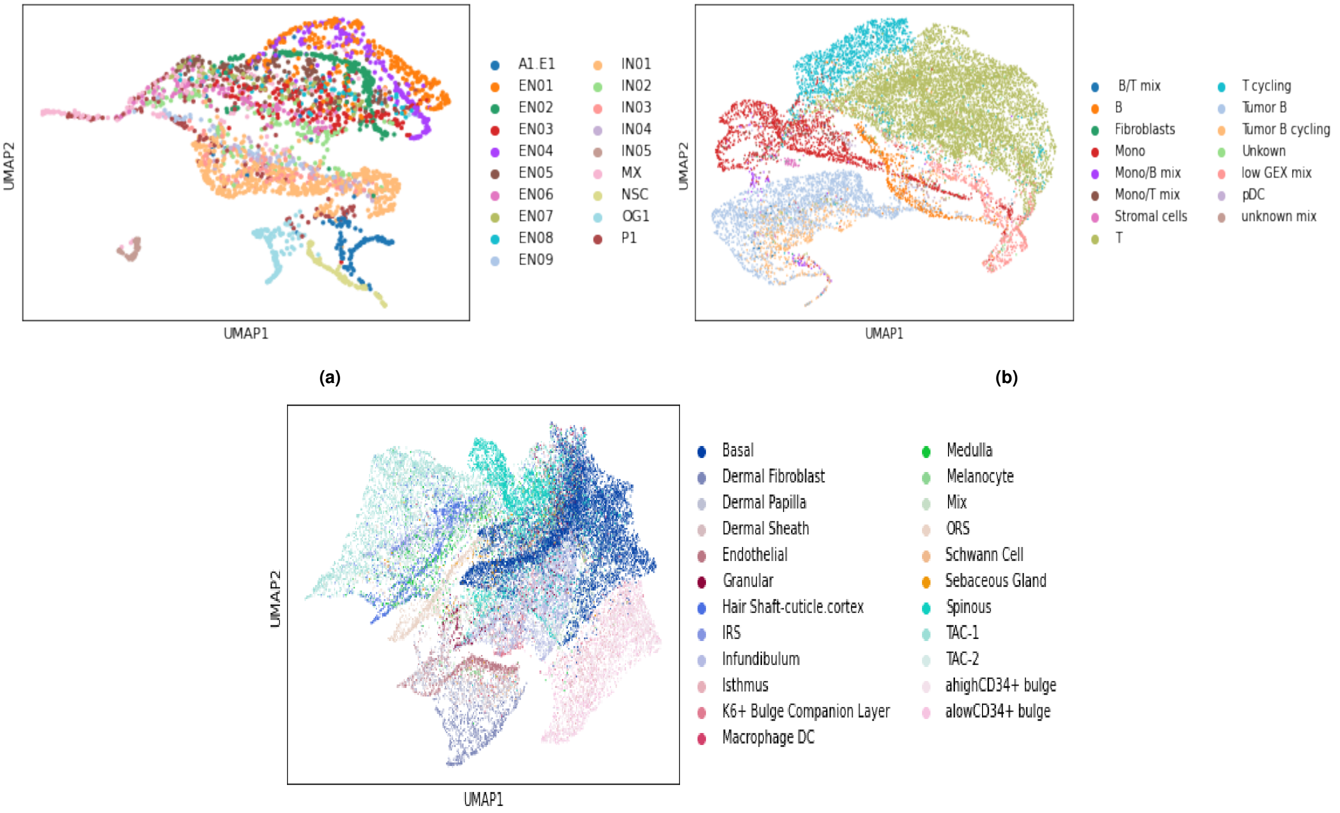


**(a)** **(b)**

**(c)**

**Fig. S4.** Umap embeddings of the reduced model considering only transcriptome data. (a) brain dataset embedding of 2781 cells from topic sapce of dimension 30. (b) B lymphoma dataset embedding 14566 cells from topic space of dimension 45. (c) skin dataset embedding of 27782 cells from topic sapce of dimension 60.

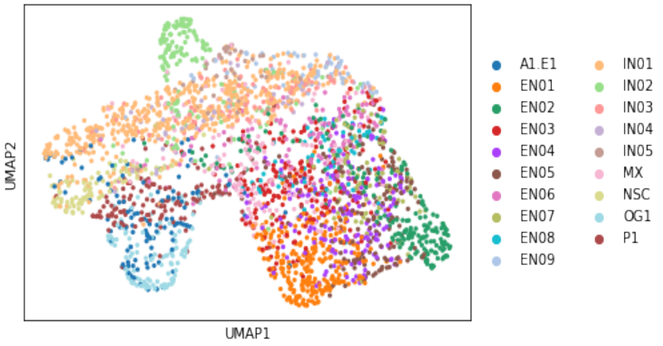


**(a)**

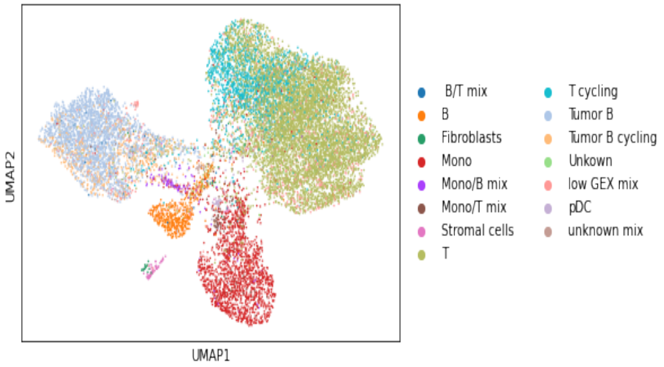


**(b)**

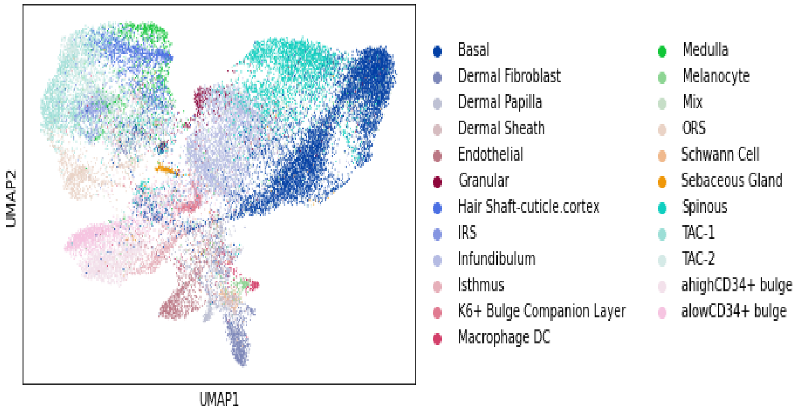


**(c)**

**Fig. S5.** Umap embeddings of the reduced model considering only chromatin accessibility data. (a) brain data set embedding of 2781 cells from topic sapce of dimension 30. (b) B lymphoma dataset embedding 14566 cells from topic space of dimension 45. (b) skin daca set embedding of 27782 cells from topic sapce of dimension 60.

| Topic | number of associated genes | number of associated regions | Biological Processes |
|---|---|---|---|
| 4 | 1716 | 8073 | synapse organization and positive regulation of cell projection organization |
| 14 | 218 | 7274 | regulation of ion transmembrane transport and syanpse organization |
| 17 | 400 | 5378 | cell junction assembly and positive regulation of nervous system development |
| 23 | 233 | 7766 | learnin memory, cognition and regulation of ion transmembrane transport |

**(a)** Brain dataset

| Topic | number of associated genes | number of associated regions | Biological Processes |
|---|---|---|---|
| 7 | 169 | 21 | regulation of cell-cell adhesion and epidermal cell differentiation |
| 46 | 138 | 1975 | positive regulation of cell cycle and chromosome organization |
| 56 | 351 | 6 | positive regulation of cell adhesion and mesenchymal cell differentiation |
| 59 | 365 | 35 | small GTPase mediated signal transduction and actin filament organization |

**(b)** Skin dataset

| Topic | number of associated genes | number of associated regions | Biological Processes |
|---|---|---|---|
| 4 | 673 | 4044 | immune response-regulating signaling pathway and positive regulation of cytokine production |
| 29 | 261 | 2972 | negative regulation of cell cycle and DNA metabolic process |
| 32 | 4583 | 2115 | immune response-activating cell surface-receptor signaling pathway and signal transduction |
| 35 | 175 | 81 | T cell differentiation and positive regulation of cytokine production |

**(c)** Lymphoma dataset

**Table S1.** Table showing the number of genes and regions associated with each topic across the three datasets. The last column annotate each topic with biological processes enriched in the associated genes

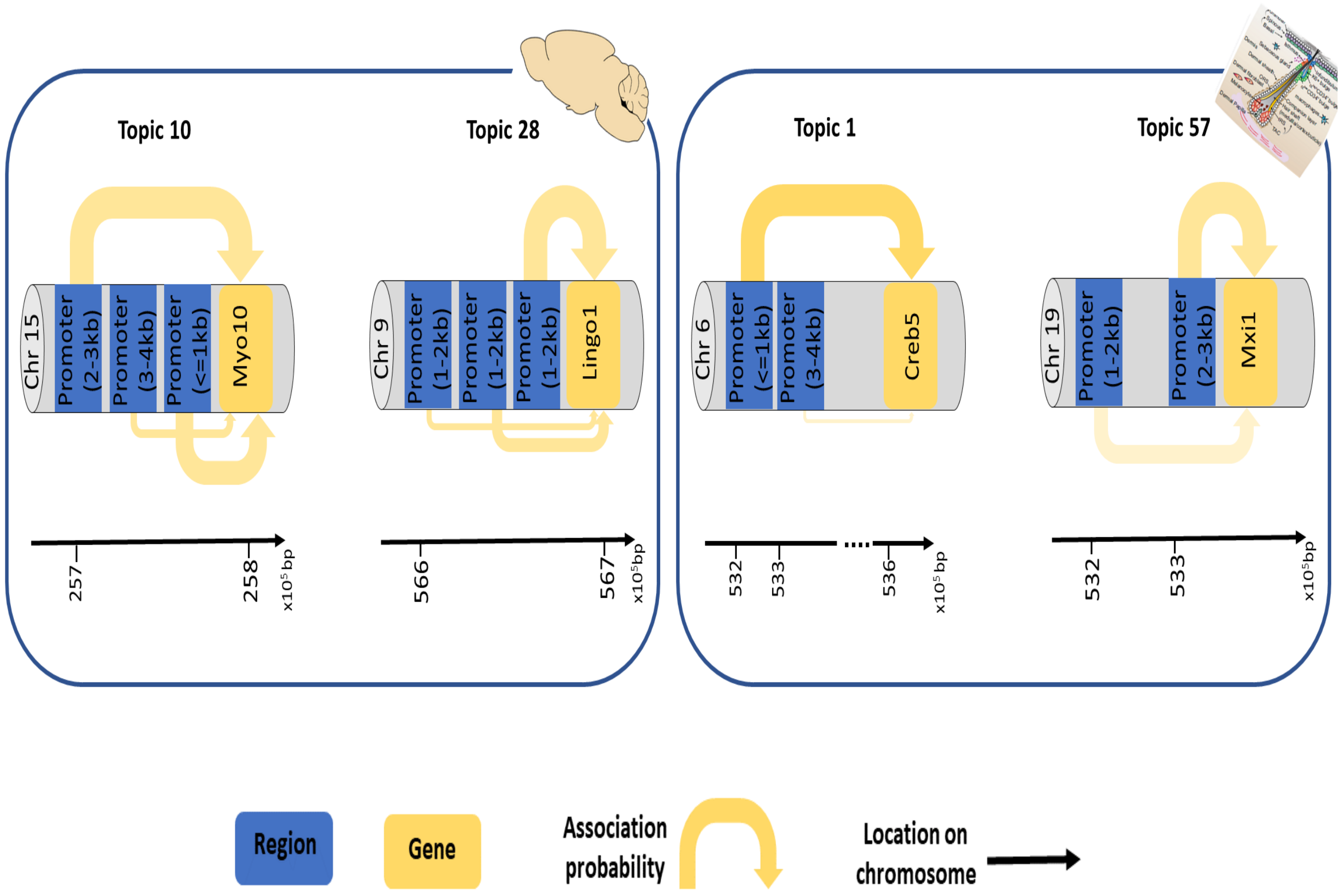


**Fig. S6.** The probabilistic interaction between genes and close promoters in a given topic(biological process). The thickness of the arrows is proportional to the interaction probability with the associated gene.
